# Ultrasound-Based Spatiotemporal Monitoring of Coagulation and Thrombolysis via Speed-of-Sound Shift Imaging

**DOI:** 10.64898/2026.06.18.733085

**Authors:** Shir Gershon, Tal Grutman, Tali Ilovitsh

## Abstract

Blood clot formation and thrombolysis are dynamic biological processes that play central roles in hemostasis, thrombosis, and thrombolytic therapy. Monitoring clot evolution is challenging, as existing approaches often rely on specialized hardware or complex acquisition protocols. This study presents dense speed-of-sound shift imaging (DSI), a noninvasive ultrasound framework for spatiotemporal monitoring of coagulation and lysis from ultrasound image sequences acquired with a single imaging transducer. DSI estimates interframe displacements using dense optical flow and reconstructs slowness-shift maps by solving a regularized inverse problem, from which relative speed-of-sound (SoS) shifts are derived. Using this approach, we quantified spatially localized acoustic signatures of material solidification, clot formation, and enzymatic clot dissolution across systems of increasing biological complexity, including thermally gelling gelatin, fibrin clots, and porcine and human whole blood. DSI detected composition-dependent clot properties, with fibrinogen primarily affecting SoS shift magnitude and thrombin primarily affecting clotting kinetics. In both porcine and human whole blood, DSI tracked the full transition from rapid clot formation to tPA-mediated thrombolysis, revealing markedly slower lysis kinetics and species-dependent differences in clotting amplitude and stabilization time. Together, these results establish DSI as a simple, ultrasound-based platform for quantitative monitoring of coagulation, thrombolysis, and related biological material transitions.

## 1. Introduction

Blood clot formation and lysis are dynamic biological processes that play critical roles in wound healing, vascular homeostasis, and therapeutic intervention. Abnormal clot formation or impaired clot breakdown can contribute to a wide range of clinical conditions, including hemorrhage, thrombosis, pulmonary embolism, and ischemic stroke ^[1–5]^. Beyond their clinical significance, coagulation and thrombolysis represent dynamic material transitions involving continuous changes in clot structure, fibrin-network organization, composition, and mechanical properties. These processes require timely evaluation because clot formation, stabilization, and therapeutic clot breakdown directly influence treatment decisions and clinical outcomes, creating a need for noninvasive methods capable of monitoring dynamic clot evolution over time.

Hemostasis involves the rapid formation of a fibrin-rich clot following vascular injury. During coagulation, thrombin-mediated fibrin polymerization produces a dynamic three-dimensional fibrin fiber network whose structure and mechanical properties depend on biochemical factors including fibrinogen concentration, thrombin activity, and calcium availability ^[6–11]^. Conversely, fibrinolysis degrades this fibrin network through plasmin-mediated clot dissolution, a process that can be therapeutically accelerated using tissue plasminogen activator (tPA) ^[12,13]^. Together, clot formation and thrombolysis represent dynamic material transitions that evolve over time and influence clot stability, treatment response, and restoration of blood flow.

However, despite the clinical importance of coagulation and thrombolysis, continuous spatiotemporal monitoring of clot evolution remains challenging. Standard coagulation assays such as activated partial thromboplastin time (aPTT) and prothrombin time (PT) provide global measurements from ex vivo plasma samples with no spatial localization and do not capture the evolving spatial structure of clots ^[14]^. Viscoelastic methods including thromboelastography (TEG) and rotational thromboelastometry (ROTEM) enable dynamic assessment of clot formation and lysis in whole blood, but remain bulk ex vivo assays that do not provide image-based spatial information on local clot heterogeneity ^[15,16]^. Alternative optical, electrochemical, microfluidic, and photoacoustic approaches have also been investigated ^[17–24]^. However, many of these methods are primarily applied in ex vivo, superficial, or controlled laboratory settings and may be less suitable for monitoring clot evolution at clinically relevant tissue depths ^[24]^. This creates a need for ultrasound-based approaches capable of noninvasive, depth-resolved monitoring of dynamic clot formation and thrombolysis.

Ultrasound is an attractive imaging modality for monitoring dynamic biological processes because it is noninvasive, portable and capable of real-time spatiotemporal imaging ^[25,26]^. Previous ultrasound studies demonstrated that coagulation alters acoustic properties including sound velocity, attenuation, and backscatter ^[27–29]^. More recent approaches, including elastography, sonorheometry, and acoustic-radiation-force-based methods, have enabled quantitative assessment of clot viscoelasticity and thrombus properties ^[30–39]^. However, many of these techniques require specialized excitation schemes, custom hardware, or indirect mechanical measurements rather than direct analysis of ultrasound data. By comparison, the feasibility of monitoring clot formation and lysis directly from conventional B-mode ultrasound data acquired with a single imaging transducer and analyzed within a standard imaging workflow remains largely unexplored. Moreover, existing ultrasound-based approaches have typically focused on either clot formation or thrombolytic breakdown, rather than providing a unified framework capable of characterizing both processes ^[40,41]^.

Here, we present dense speed-of-sound shift imaging (DSI), a noninvasive ultrasound framework for spatiotemporal monitoring of clot formation and thrombolysis using conventional B-mode image sequences acquired with a single imaging transducer. The method reconstructs dynamic speed-of-sound (SoS) shifts from interframe displacement fields estimated using dense optical flow and a regularized inversion framework. Unlike many existing coagulation-monitoring approaches, DSI operates directly on ultrasound imaging data without additional mechanical excitation or dedicated coagulation-monitoring hardware. The framework was evaluated across systems of increasing biological complexity, including thermally gelling gelatin, purified fibrin clots, porcine and human whole blood during both clot formation and tPA-mediated thrombolysis (**Figure 1**). Through these experiments, we demonstrate that DSI can provide spatially resolved information on dynamic material evolution, biochemical composition, and treatment-mediated breakdown across systems of increasing biological complexity, from gelling biomaterials to whole-blood coagulation and thrombolysis.

**Figure 1.**
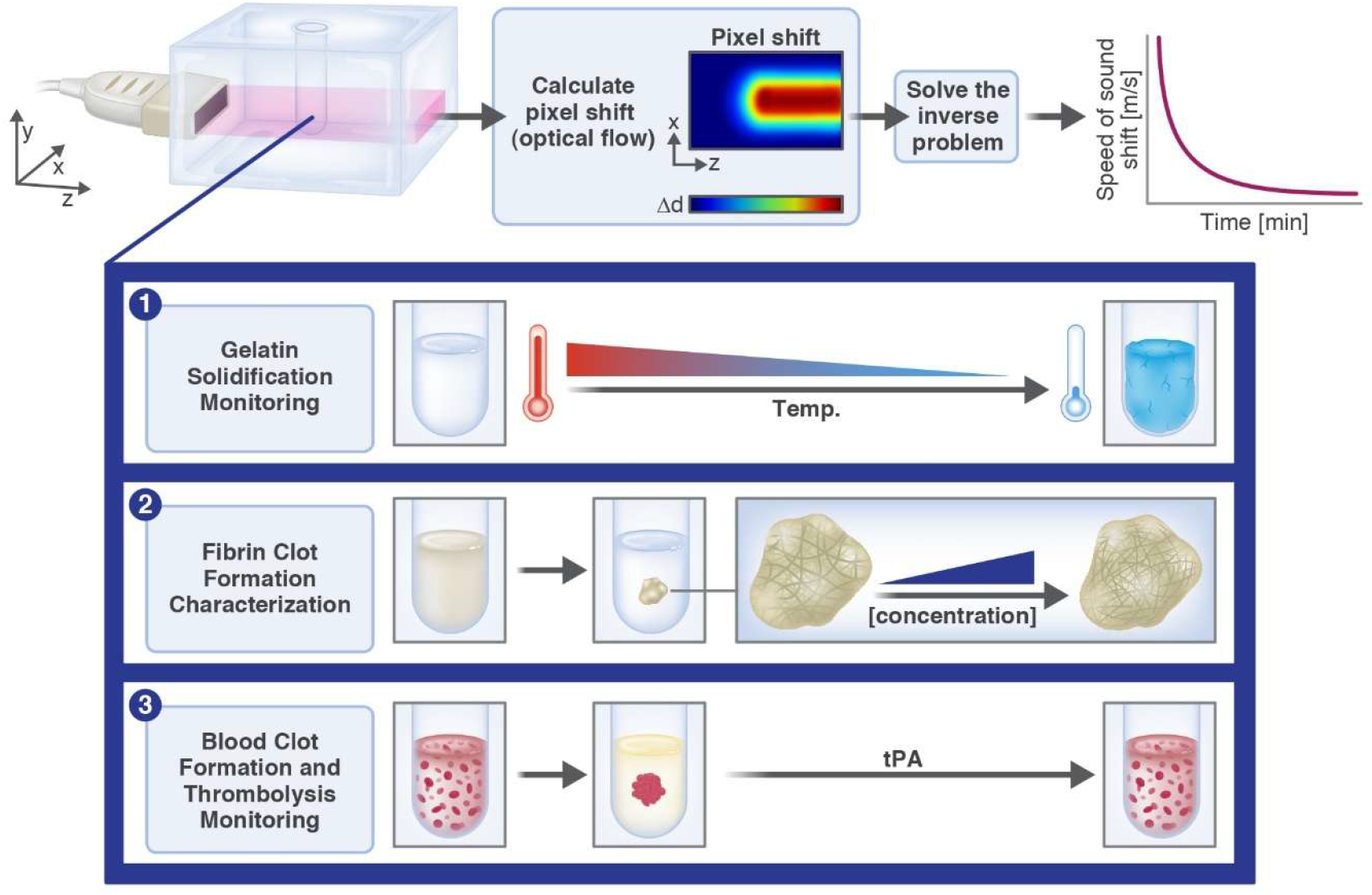
Schematic illustration of the dense speed-of-sound shift imaging (DSI) framework. Ultrasound image sequences are processed using dense optical flow to estimate interframe displacements. These measurements are incorporated into a regularized inverse model to reconstruct spatial maps of slowness shifts, from which relative speed-of-sound (SoS) changes are calculated. The resulting SoS maps and temporal profiles enable monitoring of dynamic material transitions associated with gelation, clot formation, and thrombolysis. Representative applications include: (1) monitoring the liquid-to-gel transition during temperature-induced gelatin solidification; (2) quantifying the effects of fibrinogen and thrombin concentration on clot acoustic properties; and (3) tracking coagulation and tPA-mediated clot dissolution in whole blood.

## 2. Results

### 2.1 Monitoring gelatin gelation dynamics

To provide a controlled and reproducible imaging environment, all experiments were conducted in an agarose-based tissue-mimicking phantom containing a cylindrical cavity filled with the tested material. A linear-array ultrasound transducer was positioned laterally to the phantom to image a fixed cross-sectional plane of the inclusion, and acquisitions were initiated immediately after inserting the material into the inclusion. This setup enabled repeated monitoring of the same spatial region over time, allowing characterization of the different stages of gelation, clot formation and lysis.

To evaluate whether the proposed DSI approach could capture gelation-related acoustic changes over time in a thermally gelling viscoelastic material, gelatin samples were imaged during gelation. Representative displacement maps and reconstructed slowness deviation maps from the first 20 minutes of a gelatin experiment demonstrate a spatially localized increase in values within the sample region over time, indicating progressive spatial evolution of the solidifying gelatin network (**Figure 2**A). As the slowness shifts evolved over time, the corresponding SoS shift gradually approached a stable plateau near zero.

**Figure 2.**
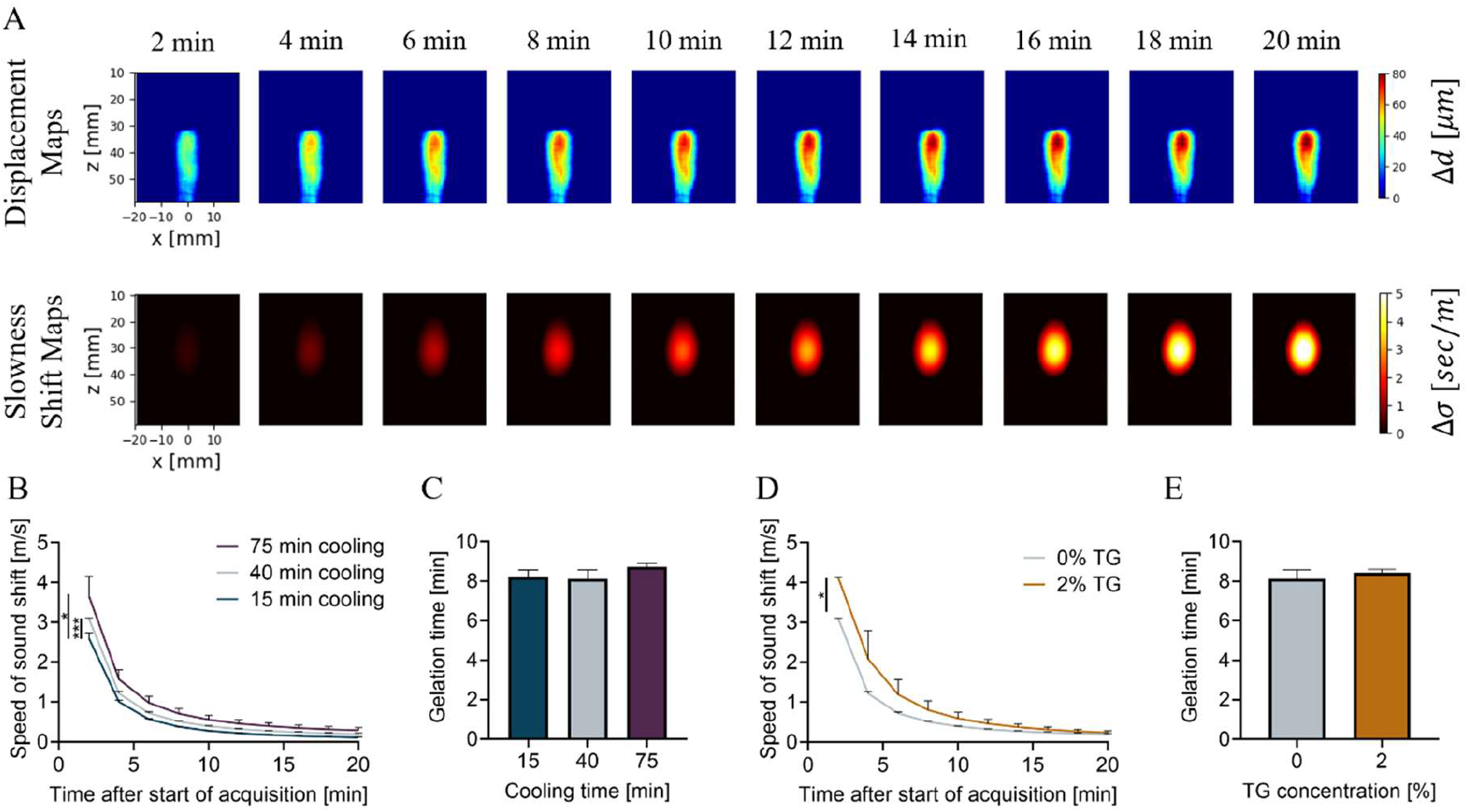
DSI assesses gelation time in gelatin samples. (A) Representative displacement maps and slowness-shift maps from the first 20 minutes of a gelatin experiment. (B) SoS shift as a function of time after the start of acquisitions in gelatin samples cooled for 15, 40, or 75 min. Cooling duration had a significant effect on SoS shift profile (Two-Way ANOVA with Tukey’s multiple comparisons test). (C) Gelation times for samples with different cooling durations, showing no significant differences (One-way ANOVA with Tukey’s multiple comparisons test). (D) SoS shift as a function of time after the start of acquisitions for samples cooled for 40 min and prepared with or without TG. TG had a significant effect on gelation kinetics (Two-Way ANOVA).(E) Gelation times for samples prepared with or without TG, showing no significant differences (unpaired t-test). Adjusted p values were \**p* < 0.05, \*\**p* < 0.01, \*\*\**p* < 0.001 and \*\*\*\**p* < 0.0001. N = 3. All data are plotted as mean ± SD.

Quantitative analysis of the SoS shift, derived from the reconstructed slowness deviation maps, showed that gelation dynamics depended on cooling history (Figure 2B). Ultrasound data acquisition was initiated for gelatin samples after cooling for 15, 40, or 75 minutes, corresponding to 53.4±0.2°C, 36.3±0.9°C, and 26.5±1.4°C, respectively. Cooling durations of gelatin solutions significantly affected the SoS shifts (p<0.01). In general, shorter cooling durations prior to imaging were associated with smaller initial SoS shifts, whereas longer cooling durations produced larger early shifts. Specifically, the initial SoS shift increased from 2.59±0.11 m s^-1^ after 15 min of cooling to 3.01±0.01 m s^-1^ after 40 min of cooling (16.2%) and to 3.53±0.41 m s^-1^ after 75 min (36.3% relative to 15 min, **Table 1**).

**Table 1.** Effect of cooling time on SoS shift and gelation time metrics (τ, 3τ, 5τ) in gelatin samples cooled for 15, 40, or 75 min before the start of ultrasound acquisitions. Values are mean ± SD (N = 3 per condition).

| Cooling time [min] | TG concentration [%] | Initial SoS shift [$\text{m s}^{-1}$] | $\tau$ [min] | $3\tau$ [min] | $5\tau$ [min] |
| --- | --- | --- | --- | --- | --- |
| 15 | 0 | 2.59±0.11 | 4.07±0.12 | 8.22±0.35 | 12.37±0.58 |
| 40 | 0 | 3.01±0.01 | 4.04±0.15 | 8.12±0.46 | 12.20±0.77 |
| 75 | 0 | 3.53±0.41 | 4.24±0.06 | 8.72±0.17 | 13.20±0.29 |

Characteristic gelation times (3τ) remained similar across cooling conditions, with values of 8.22±0.35, 8.12±0.46 and 8.72±0.17 min for the 15, 40 and 75 min cooling conditions respectively (p>0.05; Table 1). No significant differences were observed for samples with different cooling durations (Figure 2C), indicating that the cooling protocol mainly shifted the apparent starting point of gelation rather than altering the timing of the measured transition. For the 75-min cooling condition, an additional 5τ analysis was performed to estimate near-complete stabilization (∼99%), yielding a stabilization time of 13.20±0.29 minutes after the start of acquisition, corresponding to 90.20±0.29 minutes from the start of cooling. This value was within 3.01% of the stabilization time measured by the manual stirring test (93 min).

Next, to assess whether enzymatic crosslinking alters the monitored SoS dynamics during gelatin gelation, transglutaminase (TG) was added to the gelatin samples and gelation was compared with gelatin-only controls. TG is commonly used to strengthen gelatin through enzymatic crosslinking and to modify its mechanical and structural properties. TG had a significant effect on the SoS shift profile (p<0.05; Figure 2D). Specifically, adding TG increased the initial SoS shift from 3.01±0.01 m s^-1^ to 4.07±0.01 m s^-1^, corresponding to a 35.2% increase (**Table 2**). In contrast, the corresponding gelation times showed no significant differences (p>0.05; Figure 2E), increasing by 3.7% from 8.12±0.46 to 8.42±0.17 min (Table 2).

**Table 2.** Effect of TG on SoS shift and gelation time metrics (τ, 3τ) in gelatin solutions prepared with or without TG. Values are mean ± SD (N = 3 per condition).

| Cooling time [min] | TG concentration [%] | Initial SoS shift [$\text{m s}^{-1}$] | $\tau$ [min] | $3\tau$ [min] |
| --- | --- | --- | --- | --- |
| 40 | 0 | 3.01±0.01 | 4.04±0.15 | 8.12±0.46 |
| 40 | 2 | 4.07±0.01 | 4.14±0.06 | 8.42±0.17 |

### 2.2 Monitoring fibrin clot formation under varying fibrinogen and thrombin concentrations

To evaluate whether DSI could resolve biochemically driven differences in clot architecture and clotting dynamics, the framework was next applied to purified fibrin clots. To assess whether the proposed DSI approach could capture formulation-dependent changes during fibrin clot formation, fibrin clots were prepared with fibrinogen concentrations of 2.5, 3.5 or 5.0 mg mL^-1^ at a constant thrombin concentration of 0.65 U mL^-1^, or with thrombin concentrations of 0.3, 0.65 or 1.0 U mL^-1^ at a constant fibrinogen concentration of 3.5 mg mL^-1^. These ranges were selected to span normal to moderately elevated physiological fibrinogen conditions (2-4 mg mL^-1^), as well as a moderately elevated fibrinogen concentration (5 mg mL^-1^), and low to moderate thrombin concentrations relevant to fibrin clot polymerization ^[42]^.

Representative displacement maps and reconstructed slowness deviation maps from the first 2.5 minutes of a fibrin clot experiment are shown in **Figure 3**A. Both map types demonstrated a spatially localized increase in values within the sample region over time during fibrin clot formation, consistent with the spatial patterns observed during gelatin gelation.

**Figure 3.**
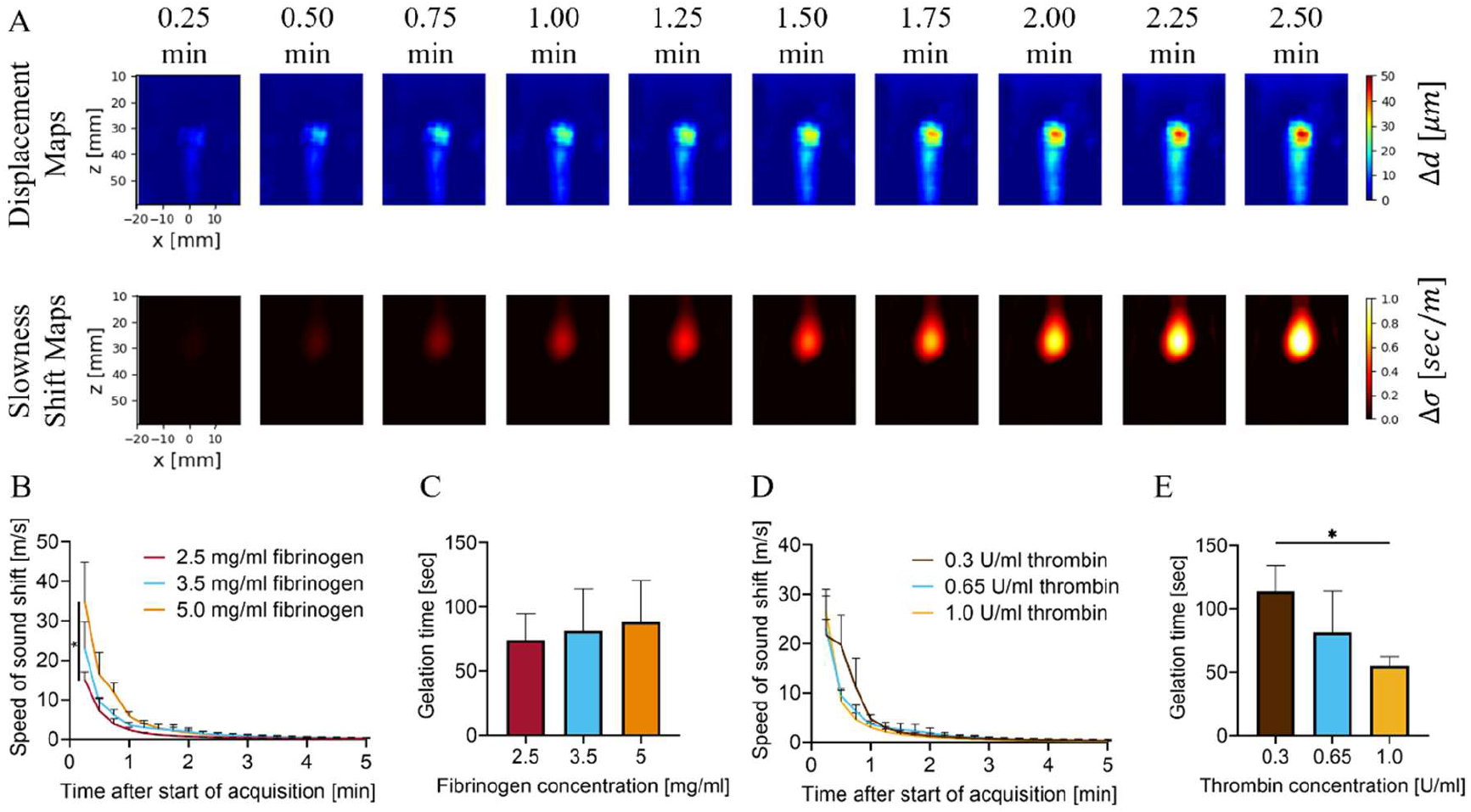
Effects of fibrinogen and thrombin concentration on SoS shift and gelation time in fibrin clots. (A)Representative displacement maps and slowness-shift maps from the first 2.5 minutes of a fibrin clot experiment. (B) SoS shift as a function of time after the start of acquisitions in fibrin clots prepared with 2.5, 3.5, or 5 mg mL^-1^ fibrinogen (0.65 U mL^-1^ thrombin and 5 mM CaCl_2_). Fibrinogen concentration had a significant effect on SoS shift (Two-Way ANOVA with Tukey’s multiple comparisons test) (C) Gelation times (3τ) for samples with different fibrinogen concentrations, showing no significant differences (1-way ANOVA with Tukey’s multiple comparison). (D) SoS shift as a function of time after the start of acquisitions in fibrin clots prepared with 0.3, 0.65 or 1.0 U mL^-1^ thrombin (3.5 mg mL^-1^ fibrinogen and 5 mM CaCl_2_). Thrombin concentration had no significant effect on SoS shift (Two-Way ANOVA with Tukey’s multiple comparisons test) (E) Gelation times for samples with different thrombin concentrations showed significant differences (1-way ANOVA with Tukey’s multiple comparisons test). Adjusted p values were \**p* < 0.05, \*\**p* < 0.01, \*\*\**p* <0.001 and \*\*\*\**p* < 0.0001. N = 3. All data are plotted as the mean ± SD.

When fibrinogen concentration was varied between 2.5, 3.5 and 5.0 mg mL^-1^ while thrombin concentration was held constant at 0.65 U mL^−1^, fibrinogen concentration significantly affected the SoS shift profile (p<0.05; Figure 3B). Increasing fibrinogen concentration produced progressively larger SoS shifts, suggesting formation of denser fibrin networks. Specifically, increasing fibrinogen concentration from 2.5 to 3.5 mg mL^−1^ increased the initial SoS shift from 15.14 ± 1.57 m s^-1^ to 23.04 ± 5.49 m s^-1^ (52.2%), while a further increase to 5.0 mg mL^−1^ increased the initial SoS shift to 34.89 ± m s^-1^ (130.4% relative to 2.5 mg mL^−1^), as summarized in **Table 3**. Although fibrinogen concentration significantly affected the initial SoS shift, the corresponding gelation times did not differ significantly across concentrations (p>0.05; Figure 3C), with gelation times of 74.20±21.78, 82.22±31.33 and 89.25±30.89 sec for 2.5, 3.5 and 5.0 mg mL^-1^ fibrinogen concentrations, respectively (Table 3). These findings indicate that increasing fibrinogen concentration primarily altered the structural properties of the developing fibrin network, leading to larger SoS shifts without substantially affecting gelation timing.

**Table 3.**
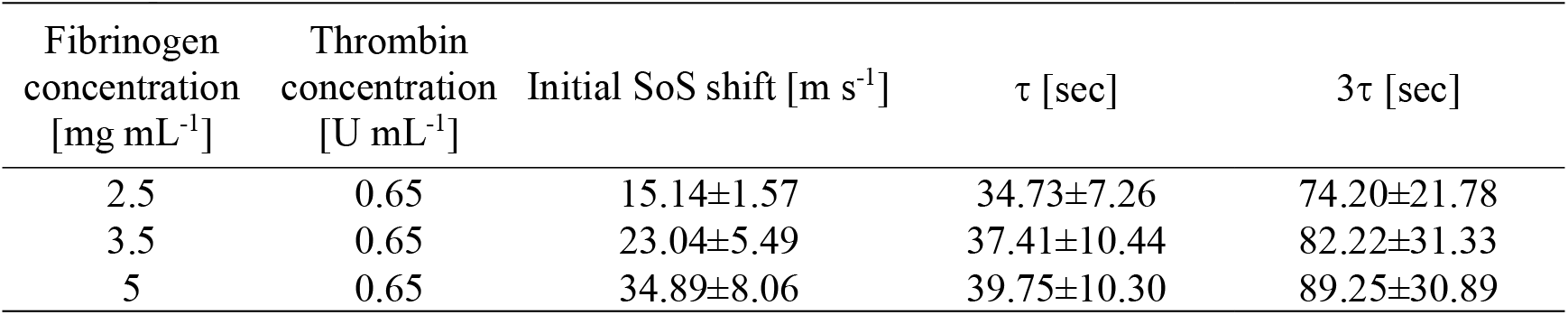
Effect of fibrinogen concentration on SoS shift and gelation time in fibrin clots. Values are mean ± SD (N = 3 per condition).

| Fibrinogen concentration [mg mL<sup>-1</sup>] | Thrombin concentration [U mL<sup>-1</sup>] | Initial SoS shift [m s<sup>-1</sup>] | τ [sec] | 3τ [sec] |
| --- | --- | --- | --- | --- |
| 2.5 | 0.65 | 15.14±1.57 | 34.73±7.26 | 74.20±21.78 |
| 3.5 | 0.65 | 23.04±5.49 | 37.41±10.44 | 82.22±31.33 |
| 5 | 0.65 | 34.89±8.06 | 39.75±10.30 | 89.25±30.89 |

When thrombin concentration was varied between 0.3, 0.65 and 1.0 U mL^-1^ while fibrinogen concentration was held constant at 3.5 mg mL^-1^, thrombin concentration had no significant effect on SoS shift profile (p>0.05; Figure 3D). The initial SoS shift increased from 21.73±2.64 m s^-1^ at 0.3 U mL^−1^ to 23.04 ± 5.49 m s^-1^ at 0.65 U mL^−1^ and 26.46±3.70 m s^-1^ at 1.0 U mL^−1^, indicating only modest differences in SoS shift magnitude across thrombin concentrations (**Table 4**). However, thrombin strongly affected clotting kinetics, with gelation times differing significantly between thrombin concentrations (p<0.05; Figure 3E). Gelation times decreased from 114.33±19.74 sec at 0.3 U mL^−1^ to 82.22±31.33 sec at 0.65 U mL^−1^ and 54.13±7.96 sec at 1.0 U mL^−1^, corresponding to decreases of 28.1% and 52.6% respectively, relative to 0.3 U mL^−1^ (Table 4). These results indicate that higher thrombin concentrations were associated with shorter gelation times, primarily accelerating clot formation while producing only modest changes in the SoS shift magnitudes.

**Table 4.** Effect of thrombin concentration on SoS shift and gelation time in fibrin clots. Values are mean ± SD (N = 3 per condition).

| Fibrinogen concentration [mg mL<sup>-1</sup>] | Thrombin concentration [U mL<sup>-1</sup>] | Initial SoS shift [m s<sup>-1</sup>] | τ [sec] | 3τ [sec] |
| --- | --- | --- | --- | --- |
| 3.5 | 0.3 | 21.73±2.64 | 48.11±6.58 | 114.33±19.74 |
| 3.5 | 0.65 | 23.04±5.49 | 37.41±10.44 | 82.22±31.33 |
| 3.5 | 1.0 | 26.46±3.70 | 28.04±2.65 | 54.13±7.96 |

Together, these results demonstrate that DSI provides complementary information regarding clot structure and clotting kinetics, enabling differentiation between factors that primarily affect fibrin-network architecture and those that primarily affect the rate of clot formation.

### 2.3 Robustness of SoS measurements to inclusion diameter

Because clot size and location may vary substantially across experimental and clinical scenarios, we evaluated whether DSI-derived measurements remained robust to variations in clot geometry. DSI reconstructs SoS changes from displacement measurements across the imaging field, therefore larger inclusions could potentially generate larger reconstructed signals even when clot composition remains unchanged. To assess the effect of clot size, fibrin clots were formed in cylindrical inclusions with diameters of 6, 8, and 10 mm while keeping biochemical clotting conditions constant: 2.5 mg mL^-1^ fibrinogen, 0.65 U mL^-1^ thrombin and 5 mM CaCl_2_.

Representative B-mode images of the three inclusion sizes demonstrate the increase in clot size with increasing inclusion diameter (Figure S1A, Supporting Information). Quantitative analysis showed a gradual increase in SoS shift magnitude caused by increasing inclusion diameter. However, this effect did not reach statistical significance (p>0.05; Figure S1B, Supporting Information). Consistent with this trend, the initial SoS shift tended to increase with inclusion diameter, from 13.70±0.37 m s^-1^ at 6mm inclusions to 15.14±1.57 m s^-1^ at 8mm inclusions and 18.17±3.98 m s^-1^ at 10mm inclusions (Table S1, Supporting Information). Similarly, the corresponding gelation times remained comparable across inclusion diameters, with values of 66.17±10.85, 74.20±21.78 and 85.23±12.16 s for 6, 8, and 10 mm inclusions, respectively, with no significant differences between groups (p>0.05; Figure S1C and Table S1, Supporting Information).

### 2.4 Robustness of fibrin clot SoS measurements to inclusion placement

One of the principal advantages of ultrasound is its ability to interrogate structures at clinically relevant tissue depths. Therefore, for DSI to be useful across different imaging scenarios, its measurements should remain stable across variations in clot location within the imaging field. To evaluate the effect of inclusion placement, fibrin clots were formed in 8-mm diameter cylindrical inclusions positioned at different axial and lateral locations relative to the transducer: 1.5 cm depth, 3.0 cm depth (reference), 4.5 cm depth, and 3.0 cm depth with a 1.0 cm lateral shift, while maintaining the same biochemical clotting conditions: 2.5 mg mL^-1^ fibrinogen, 0.65 U mL^-1^ thrombin and 5 mM CaCl_2_.

Representative B-mode images of the four inclusion placements are shown in Figure S2A (Supporting Information). Inclusion position had no significant effect on the SoS shift (p>0.05; Figure S2B, Supporting Information). Initial SoS shifts remained comparable across positions, with values of 18.34±2.33, 15.14±1.57, 16.80±4.46, and 15.28±2.69 m s^-1^ for inclusions positioned at 1.5 cm depth, 3.0 cm depth, 4.5 cm depth and 3.0 depth with a 1.0 cm lateral shift, respectively (Table S2, Supporting Information). Similarly, gelation times remained comparable across positions, with values of 81.22±27.59, 74.20±21.78, 88.24±20.04, and 70.18±7.58 s, respectively, with no significant differences between groups (p>0.05; Figure S2C and Table S2, Supporting Information). Together, these results demonstrate that DSI-derived SoS shift measurements and gelation times remained stable across the tested range of imaging depths and lateral positions.

### 2.5 Monitoring porcine blood clot formation and thrombolysis

To evaluate whether DSI could monitor treatment-mediated clot breakdown, thrombolysis was initiated using tPA following clot stabilization.

Porcine blood clotting produced a SoS shift trajectory comparable to that observed during fibrin clot formation (**Figure 4**A), with no significant differences in the SoS shift profiles between groups (p>0.05). The initial SoS shift was 11.29±2.46 m s^-1^ in porcine blood and 15.14±1.57 m s^-1^ in fibrin clots. These results support the relevance of our fibrin clot model for subsequent whole blood experiments.

**Figure 4.**
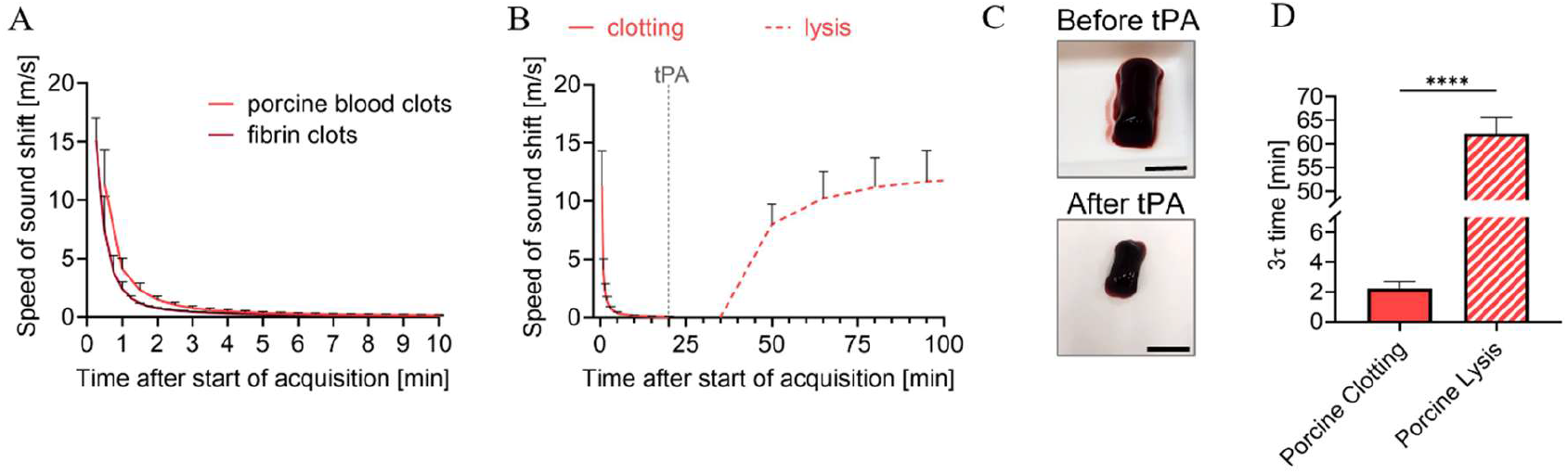
Monitoring porcine blood clot formation and thrombolysis. Porcine citrated whole blood was recalcified with 15 mM CaCl2 to initiate clotting. 6 µg mL-1 tPA was added after 20 min for thrombolysis experiments. (A) SoS shift as a function of time during porcine whole-blood clotting compared with fibrin clotting, showing qualitatively similar clotting trajectories (Two-Way ANOVA with Tukey’s multiple comparisons test). (B) SoS shift during clotting and subsequent lysis, with the dashed line indicating the time of tPA addition. (C) Representative images of porcine clots after clot formation and after tPA-mediated lysis. Black scale bars = 1 cm. (D) Comparison of 3τ times for clotting and lysis, showing that lysis occurred on a substantially slower timescale than clot formation (unpaired two-tailed t-test). Adjusted p values were \**p* < 0.05, \*\**p* < 0.01, \*\*\**p* < 0.001 and \*\*\*\**p* < 0.0001. N = 3. All data are plotted as the mean ± SD.

To evaluate whether the method could also capture clot breakdown, tPA was added after 20 min of clotting, and the resulting SoS shift was monitored over time. Following tPA addition, the SoS shift gradually increased toward a new plateau, reaching a SoS shift of 12.88±2.48 m s^-1^, comparable to the initial SoS shift observed during clot formation. The full lysis time course is shown in Figure 4B. This tPA-mediated clot breakdown was also visibly evident in photographs of the resulting clots, which show a visible reduction in clot size after treatment (Figure 4C).

The thrombolysis-related acoustic response was substantially slower than clot formation (p<0.0001). This difference in kinetics was also reflected in the corresponding lysis time, with lysis requiring longer times than clotting (Figure 4D). 3τ increased from 135.35±26.06 s (2.26±0.43 min) during clotting to 62.16±3.47 min during lysis.

### 2.6 Monitoring human blood clot formation and thrombolysis

To assess the translational relevance of DSI under clinically relevant coagulation conditions, the framework was next evaluated in human whole blood during both clot formation and tPA-mediated thrombolysis. Following clot stabilization, thrombolysis was initiated by addition of tPA.

Representative B-mode images and reconstructed slowness deviation maps from a human blood clot thrombolysis experiment are shown in **Figure 5**A. The slowness maps demonstrated a spatially localized decrease in values within the clot region over time during thrombolysis. This behavior was qualitatively consistent with the corresponding B-mode images, which showed a progressive reduction in the visible clot area over time.

**Figure 5.**
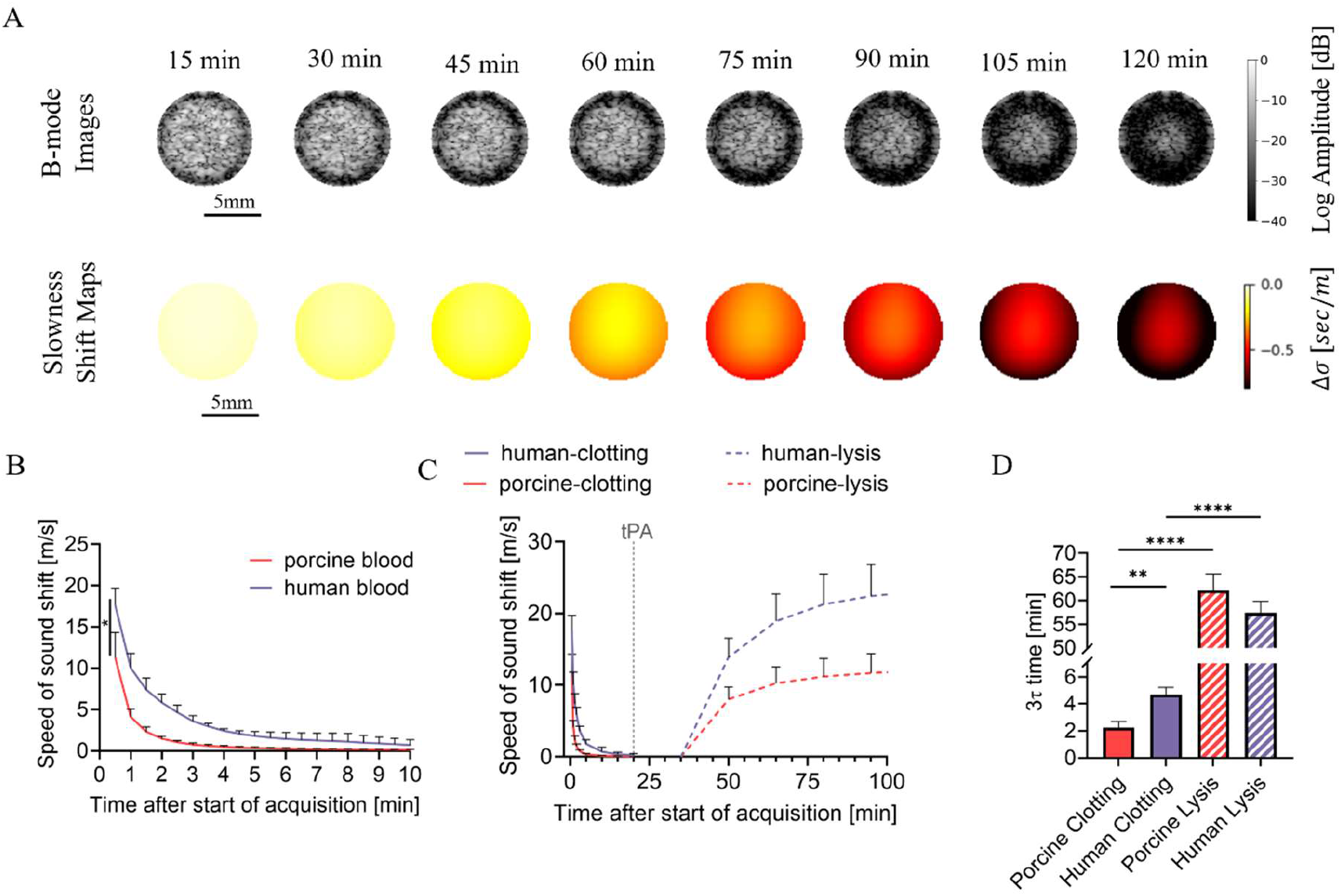
Monitoring human whole-blood clot formation and thrombolysis. Citrated human whole blood was recalcified with 15 mM CaCl2 to initiate clotting. 6 µg mL-1 tPA was added after 20 min for thrombolysis experiments. (A) Representative B-mode images and slowness-shift maps during the first 2 hours of human blood clot thrombolysis. The slowness-shift maps demonstrate a spatially localized decrease in values within the clot region over time during thrombolysis. (B) SoS shift as a function of time during human whole-blood clotting compared with porcine whole-blood clotting, showing qualitatively similar clotting trajectories, with significantly larger SoS shifts and slower stabilization observed in human blood compared with porcine blood (Two-Way ANOVA). (C) SoS shift during clotting and subsequent lysis in porcine and human blood samples, with the dashed line indicating the time of tPA addition. (D) Comparison of 3τ times for clotting and lysis in porcine and human blood, showing that lysis occurred on a substantially slower timescale than clot formation in both species. Clotting times are significantly faster in porcine blood compared with human blood, whereas thrombolysis times remained comparable between species (unpaired two-tailed t-tests). Adjusted p values were *p < 0.05, and **p < 0.01, ***p < 0.001 and ****p < 0.0001. N = 3. All data are plotted as the mean ± SD.

Human and porcine blood exhibited similar clotting-associated SoS shift trajectories following recalcification (Figure 5B). However, human blood exhibited significantly larger SoS shifts (p<0.05) and longer clotting times (p<0.01) compared with porcine blood. The initial SoS shift was 11.29±2.46 m s^-1^ in porcine blood and 17.64±1.68 m s^-1^ in human blood. Both porcine and human blood clots exhibited gradual SoS shift evolution during thrombolysis, with substantially slower dynamics compared with clot formation. In the resulting lysis trajectories, the SoS shift gradually increased toward a new plateau, reaching a SoS shift of 23.45±4.58 m s^-1^, comparable to the initial SoS shift observed during clot formation (Figure 5C).

The extracted clotting times increased from 135.35±26.06 s (2.26±0.43 min) for porcine blood to 281.44±31.11 s (4.69±0.52 min) for human blood (Figure 5D). Despite these differences, both blood models reached clot stabilization well before tPA administration at 20 min, supporting the use of a shared thrombolysis timeline for subsequent experiments. Quantitative analysis further showed that thrombolysis in human blood occurred on a substantially slower timescale than clot formation (p<0.0001). 3τ increased from 281.44±31.11 s (4.69±0.52 min) during clotting to 57.39±2.42 min during lysis. The extracted lysis times were comparable between porcine blood and human blood (p>0.05), measuring 62.16±3.47 min in porcine blood and 57.39±2.42 min for human blood. A summary of the extracted SoS shift values and corresponding clotting and lysis times across fibrin, porcine whole blood and human whole blood is provided in **Table 5**. Representative photographs of porcine and human blood clots before and after tPA-mediated thrombolysis are provided in Figure S3 (Supporting Information).

**Table 5.** Comparison of clotting and thrombolysis parameters across fibrin, porcine whole blood and human whole blood. Values are mean ± SD (N = 3 per condition).

| Model | Initial SoS shift (clotting) [$\text{m s}^{-1}$] | Final SoS shift (lysis) [$\text{m s}^{-1}$] | $3\tau$ clotting [min] | $3\tau$ lysis [min] |
| --- | --- | --- | --- | --- |
| Fibrin | $15.14 \pm 1.57$ | N/A | $1.24 \pm 0.36$ | N/A |
| Porcine | $11.29 \pm 2.46$ | $12.88 \pm 2.48$ | $2.26 \pm 0.43$ | $62.16 \pm 3.47$ |
| Human | $17.64 \pm 1.68$ | $23.45 \pm 4.58$ | $4.69 \pm 0.52$ | $57.39 \pm 2.42$ |

Together, these results demonstrate that DSI can monitor both clot formation and tPA-mediated clot breakdown in whole blood, while capturing both the kinetic asymmetry between rapid clotting and slower thrombolysis and the species-dependent differences in SoS shift magnitude and clotting kinetics.

## 3. Discussion

This study demonstrates that DSI enables noninvasive spatiotemporal monitoring of coagulation and thrombolysis directly from conventional B-mode ultrasound data. The present work extends previous DSI implementations developed for monitoring thermally induced acoustic changes during thermal ablation and cryoablation ^[43,44]^ toward intrinsic material processes associated with clot formation, structural maturation, and enzymatic clot breakdown. Across progressively more complex systems, from gelatin gelation to whole-blood thrombolysis, the method remained sensitive to dynamic material transitions associated with clot formation, structural maturation, and enzymatic clot breakdown. Importantly, these measurements were obtained using a single imaging transducer without specialized excitation schemes, suggesting a potential path toward clinically accessible ultrasound-based coagulation monitoring.

One of the central findings of this study is that DSI differentiated between structural and kinetic determinants of clot formation. Increasing fibrinogen concentration primarily increased the magnitude of the SoS response, whereas increasing thrombin concentration predominantly accelerated clotting kinetics with comparatively smaller effects on SoS shift magnitude. These findings are consistent with previous reports showing that elevated fibrinogen concentrations produce denser fibrin networks with larger fiber diameters, reduced pore size, and increased turbidity, while thrombin primarily regulates polymerization rate and clotting lag time ^[11]^. Accordingly, fibrinogen concentration is expected to alter acoustic wave propagation through changes in fibrin-network structure, whereas thrombin concentration primarily affects the rate of the fluid-to-solid transition.

Together, these results demonstrate that DSI captures complementary aspects of clot formation beyond a single endpoint measurement, with the reconstructed SoS shifts potentially reflecting changes in clot density, network organization, viscoelasticity, and polymerization dynamics. The clotting timescales observed in the present fibrin experiments were also consistent with previous fibrin-clot studies reporting polymerization dynamics on the order of seconds to minutes depending on fibrinogen and thrombin concentration ^[11,45]^, further supporting the physiological relevance of the DSI-derived gelation estimates.

The gelatin experiments further support the interpretation that DSI is sensitive to evolving material state and network formation during dynamic solidification. The measured SoS shifts during gelatin gelation, which were in the range of several meters per second, are consistent with values reported in previous ultrasound studies of gelatin gelation ^[46,47]^. The observed positive SoS shifts are consistent with the transition from a liquid-like material to a progressively more connected gel network, which is expected to alter acoustic wave propagation through the gelatin, based on changes in density, compressibility, and network structure ^[46,47]^. Cooling history substantially altered the measured SoS trajectories, consistent with previous studies demonstrating that gelatin gelation is strongly path-dependent and influenced by thermal history and quench conditions ^[46,48]^. Similarly, enzymatic crosslinking with TG increased the magnitude of the measured SoS shifts without substantially altering gelation timing, consistent with prior reports showing that TG strengthens gelatin network structure while largely preserving bulk gelation behavior ^[49,50]^. Together, these findings suggest that DSI captures both the temporal evolution and structural state of dynamically solidifying biomaterials.

Importantly, the framework remained sensitive to clot formation and thrombolysis in whole blood despite the substantially greater biological complexity of cellular and plasma-containing systems. The observed clotting-associated SoS shifts and clotting timescales were within the range reported in prior ultrasound studies of coagulation and sonorheometry in whole blood ^[27,38,51,52]^. Human blood exhibited larger SoS shifts and slower clotting kinetics compared with porcine blood, consistent with previous reports describing species-dependent differences in fibrinogen contribution, clot propagation, clot density and structure, and overall coagulation dynamics ^[53–55]^. Notably, DSI also resolved the pronounced kinetic asymmetry between rapid coagulation and substantially slower tPA-mediated thrombolysis. This observation is biologically plausible because fibrin polymerization occurs rapidly during early clot formation, whereas fibrinolysis depends on progressive enzymatic penetration and degradation of an already established fibrin network ^[56,57]^. During thrombolysis, progressive degradation of the fibrin network is expected to reduce the acoustic differences between the clot and the surrounding fluid phase. The observed evolution toward a stable post-lysis plateau is therefore consistent with gradual structural breakdown of the clot and stabilization of the post-lysis state. The ability to monitor both clot formation and treatment-mediated clot breakdown may therefore be particularly relevant for longitudinal monitoring of thrombolytic response.

Beyond biochemical composition, the geometry and placement experiments provided insight into the robustness of the proposed framework. Although a modest diameter-dependent trend was observed in SoS shift magnitude, this effect did not reach statistical significance over the tested diameter range, and inclusion diameter did not significantly affect the extracted gelation times. Importantly, DSI-derived measurements remained stable across inclusion depths ranging from 1.5 to 4.5 cm and across different lateral positions, supporting the robustness of the approach across clinically relevant imaging geometries (Figures S1–S2). Together, the geometry and placement experiments suggest that DSI measurements are primarily driven by clot composition and evolution rather than by moderate variations in clot geometry or imaging location within the field of view.

Although the present results demonstrate clear sensitivity of DSI to clot evolution, the precise biophysical origins of the measured SoS shifts likely involve multiple coupled processes, including changes in compressibility, density, fibrin network organization, viscoelasticity, and cellular redistribution during coagulation and lysis. Future studies combining DSI with direct structural and mechanical characterization will therefore be important for further resolving the physical determinants of the observed acoustic responses. Future work should also investigate real-time implementation of the DSI framework to further support clinical integration.

Several limitations should also be considered for future clinical translation. The current experiments were performed in static phantom configurations and therefore did not incorporate physiological flow, vessel pulsatility, respiratory motion, or tissue motion, all of which may influence displacement estimation in vivo. Future studies should therefore evaluate DSI under hemodynamic flow conditions and integrate motion-compensation strategies for vascular imaging applications ^[58]^. In addition, the simplified cylindrical clot geometries used here do not fully capture the structural heterogeneity and partial occlusion patterns observed in native thrombi ^[59,60]^. Further work will also be needed to evaluate DSI during clinically relevant thrombolysis paradigms, including ultrasound-enhanced thrombolysis and sonothrombolysis approaches ^[61,62]^, and to characterize the sensitivity of the framework across varying tPA concentrations and treatment conditions. Beyond coagulation and thrombolysis, the demonstrated sensitivity of DSI to dynamic material transitions suggests potential applications in monitoring the curing, gelation, or degradation of hemostatic materials and biomaterials used in wound care, tissue engineering, and regenerative medicine, including hydrogels, tissue adhesives, and clot-promoting materials designed to modify clot formation or stability ^[63]^.

The present results establish DSI as a practical ultrasound-based framework for monitoring dynamic clot evolution and thrombolytic response using conventional B-mode ultrasound acquisitions. Across all tested systems, DSI successfully identified dynamic material transitions associated with gelation, fibrin polymerization, whole-blood clot formation, and enzymatic clot breakdown, demonstrating its ability to monitor multiple stages of clot evolution. By operating directly on ultrasound imaging data acquired with a single imaging transducer, DSI may provide a practical and clinically accessible approach for integrating spatiotemporal coagulation monitoring into existing ultrasound workflows.

## 4. Conclusion

This study demonstrates that DSI can monitor dynamic acoustic changes during gelation, clot formation and lysis using conventional B-mode ultrasound data acquired with a single imaging transducer. Across gelatin, purified fibrin clots, and whole blood, the method captured spatially localized and temporally evolving responses that were sensitive to biochemical composition and thrombolytic breakdown, while remaining robust to variations in clot geometry and placement. Together, these findings establish DSI as a simple ultrasound-based framework for noninvasive monitoring of clot-related material dynamics. In this context, DSI may complement existing ultrasound-based coagulation and thrombus-characterization methods by extracting spatially resolved acoustic information directly from standard imaging data, thereby providing a foundation for future translation toward treatment monitoring and evaluation of hemostatic materials.

## 5. Methods

### 5.1. Materials and sample preparation

#### 5.1.1. Tissue-Mimicking Phantom Preparation

A tissue mimicking phantom was prepared by dissolving 4.5 grams of agarose powder (A10752.36, Thermo Fisher Scientific, MA, USA) in 300 ml of distilled water and heating the mixture in a microwave until boiling. The resulting agarose solution was left to cool at room temperature for 10 minutes and was then stirred with a magnetic stirrer at 300 rpm for an additional 10 minutes (Cimarec+™ Stirring Hotplates Series, SP88857108, Thermo Fisher Scientific, Waltham, MA, USA). 0.7 grams of silicon carbide powder (357391, Sigma Aldrich, MO, USA) were added to the solution as acoustic scatterers and the mixture was stirred for 30 additional minutes to ensure homogeneous particle distribution. The mixture was poured into a mold (9 cm × 6 cm × 3 cm) containing a cylindrical rod.

Once the agarose phantom had fully gelled, it was carefully removed from the mold, leaving a cylindrical hole in the phantom.

The cylindrical inclusion of the agarose phantom measured 25 mm in height and, unless otherwise specified, 8 mm in diameter, and was positioned 3 cm from the transducer, centered laterally in the imaging plane.

#### 5.1.2 Gelatin Solution Preparation

Gelatin was used as a thermally gelling viscoelastic material to validate the proposed approach. A 12% (w v^-1^) gelatin solution was prepared by dissolving 12 grams of gelatin powder (G9382, Type B, 225 Bloom, bovine skin, Sigma Aldrich, MO, USA) in 100 ml of distilled water. The solution was heated in a microwave until fully dissolved and then cooled at room temperature for 15, 40, or 75 minutes prior to imaging.

Gelatin temperature was measured immediately before imaging using a digital probe thermometer (TP301, Yacumama). The gelatin solution was then poured into the cylindrical void of the pre-solidified agarose phantom, and ultrasound imaging was initiated immediately.

For independent validation of gelation timing in gelatin samples, a manual magnetic stirring test was performed at 300 rpm (Cimarec+™ Stirring Hotplates Series, SP88857108, Thermo Fisher Scientific, Waltham, MA, USA). The gelation endpoint was defined as the time point at which visible stirring-induced motion of the magnetic rod ceased.

#### 5.1.3 Gelatin-Transglutaminase Preparation

Gelatin solutions containing TG were prepared to evaluate the effect of enzymatic crosslinking on the monitored dynamics, as TG crosslinking has been shown to alter the rheological and structural properties of gelatin while preserving its sol-gel transition behavior ^[50]^. A TG stock solution was prepared by heating 10 ml of distilled water in an ultrasonic bath (Elmasonic P 60H, Elma Schmidbauer GmbH, Singen, Germany) to 50°C for 3 minutes. 2 grams of TG (Ajinomoto ACTIVA WM, catalog no. 20210216, ENCO, Israel) were gradually added to the heated water while mixing with a vortex mixer (36110740, Gilson, WI, USA) at medium speed to minimize clumping. The solution was further sonicated at 50°C, 37 kHz, and 100% power for 7 minutes to ensure complete dissolution.

The TG stock solution was added to a partially cooled gelatin solution 15 minutes after the start of gelatin cooling. To obtain a final composition of 12% (w v^-1^) gelatin and 2% (w v^-1^) TG, 12 gr gelatin powder was dissolved in 90 ml distilled water, after which the TG stock solution was added. The resulting gelatin-TG solution was mixed with a magnetic stirrer for 2 minutes at 200 rpm to achieve homogeneity. The gelatin-TG solution was poured into the cylindrical void of the pre-solidified agarose phantom after 40 minutes of total gelatin cooling, allowing 25 minutes of interactions between the gelatin and the TG prior to the start of ultrasound acquisitions.

#### 5.1.4 Thrombin and fibrinogen stock solutions and clot formulation

Thrombin (605157, Sigma-Aldrich, MO, USA) was reconstituted in 2.5 ml DPBS (−/−) (02-023-1A-24, Sartorius, Göttingen, Germany) to prepare a 400 U ml^-1^ stock solutions. For preparation of the thrombin working solution, 0.125 ml of the 400 U ml^-1^ stock was diluted into 50 ml DPBS (−/−) to obtain a final concentration of 1 U ml^-1^. Fibrinogen (341573, Sigma-Aldrich, MO, USA) was dissolved in DPBS (−/−) to prepare a 20 mg mL^-1^ stock (1 g in 50 mL). Thrombin and fibrinogen stock and working solutions were aliquoted and stored at −20°C until use.

On the day of imaging, fibrinogen and thrombin aliquots were thawed and mixed immediately before acquisition. All mixtures were prepared in DPBS (−/−) supplemented with calcium chloride (CaCl_2_) with a final concentration of 5 mM, consistent with previously reported in vitro fibrin polymerization conditions ^[11,42]^, to achieve the desired final fibrinogen and thrombin concentrations. Ultrasound imaging was initiated immediately after mixing.

To assess the effect of fibrinogen concentration, thrombin concentration was held constant at 0.65 U ml^-1^ while fibrinogen was varied between 2.5, 3.5, and 5 mg mL^-1^. Conversely, to examine the effect of thrombin concentration, fibrinogen concentration was held constant at 3.5 mg mL^-1^ while thrombin was varied between 0.3, 0.65, and 1 U m^-1^. The selected fibrinogen concentrations span the normal physiological range in humans (2-4 mg mL^-1^), as well as elevated fibrinogen conditions, while the thrombin concentrations were selected to cover low to moderate clotting conditions ^[42]^.

In additional fibrin experiments examining the effect of cylindrical inclusion position and inclusion diameter size on SoS shift and gelation time, fibrin clots were prepared using 2.5 mg mL^-1^ fibrinogen and 0.65 U mL^-1^ thrombin, and 5 mM CaCl_2_. For diameter-variation experiments, inclusions with diameters of 6 mm and 10 mm were tested and compared to the default 8 mm inclusion. For position-variation experiments, inclusions were placed at depths of 1.5 cm and 4.5 cm, as well as at a depth of 3 cm with a 1 cm lateral shift and compared to the default placement configuration of 3 cm depth and laterally centered.

#### 5.1.5 Porcine blood clotting and tPA-mediated thrombolysis

Porcine whole blood was used to assess the applicability of the proposed method for monitoring clot formation and lysis in a physiologically relevant blood model. Blood was obtained from Lahav CRO Comprehensive Preclinical Services (Institute of Animal Research, Israel) in collection tubes containing 3.2% sodium citrate at a standard 9:1 blood-to-anticoagulant ratio. Upon arrival, porcine whole blood was stored in the original collection tubes at 4°C until use. All porcine blood experiments were performed within two weeks of blood arrival.

Coagulation was initiated immediately before acquisition by addition of CaCl_2_ to 1 mL of citrated blood, resulting in a final CaCl_2_ concentration of 15 mM, consistent with previously reported ultrasound-based blood clotting and lysis studies ^[22,38,40]^. For clotting-only experiments, samples were monitored for 90 minutes, with ultrasound imaging initiated immediately after the start of clotting.

For thrombolysis experiments, tPA (T-PA Protein, Human [HEK293, His], HY-P71051, MedChemExpress, Monmouth Junction, NJ, USA) was reconstituted in double-distilled water, aliquoted, and stored at −20 °C until use. Prior to the experiments, tPA aliquots were thawed and diluted in DPBS (−/−) to a concentration of 6 µg mL^-1^. This concentration was selected as a value above the clinically motivated 3.15 µg mL^-1^ benchmark used in human-blood-based studies, given previous reports that porcine clots are relatively resistant to tPA-mediated thrombolysis and may require higher thrombolytic concentrations to achieve measurable clot breakdown in vitro ^[64,65]^. For thrombolysis experiments, blood was first allowed to clot for 20 min, after which the tPA solution was added. This timepoint was selected based on the measured clotting kinetics, to ensure that clot formation had largely stabilized before thrombolysis was initiated. Ultrasound imaging was initiated immediately after the addition of the tPA solution.

#### 5.1.6 Human blood clotting and thrombolysis

Human whole blood was used to evaluate the applicability of the DSI framework in a clinically relevant blood model. Human whole blood experiments were approved by the Tel Aviv University Institutional Ethics Committee (approval no. 0011538-1) and conducted in accordance with institutional guidelines. Blood samples were obtained as citrated whole blood (3.2% sodium citrate) from the blood bank at Sheba Medical Center from a healthy adult donor, after written informed consent. Upon arrival, the blood was divided into aliquots and stored at 4°C until use. All human blood experiments were performed within two weeks of blood arrival.

Prior to imaging, coagulation was initiated immediately before acquisition by addition of CaCl_2_ to 1 ml of citrated blood, resulting in a final CaCl_2_ concentration of 15 mM. For clotting-only experiments, samples were monitored for 90 minutes, with ultrasound imaging initiated immediately after the start of clotting.

For thrombolysis experiments, tPA was prepared and applied as described above. Although lower tPA concentrations are commonly used as clinically motivated benchmarks in human-blood thrombolysis studies, a concentration of 6 µg mL^-1^ was chosen for the human clot experiments, to allow direct comparison to porcine blood clot experiments under matched experimental conditions.

Blood was allowed to clot for 20 min prior to tPA addition, a time point selected to ensure that clot formation had largely stabilized relative to the measured gelation times and to enable direct comparison between human and porcine blood under matched thrombolytic conditions.

### 5.2. Ultrasound imaging

Ultrasound imaging was performed using an L12-5 50 mm linear-array transducer (ATL Philips, WA, USA) placed in contact with the lateral surface of the agarose phantom. The transducer was oriented to image a cross-sectional plane at a fixed phantom height, such that the cylindrical inclusion was visualized in cross-section. Prior to each experiment, the agarose phantom was allowed to stabilize at room temperature. To minimize reflections, a 6 mm rubber layer was placed around the phantom and coupled with ultrasound gel. A programmable ultrasound system (Vantage 256, Verasonics Inc., WA, USA) was used to image a 59 mm × 40 mm field of view. A plane-wave steering acquisition protocol was used to acquire nine plane waves per imaging sequence. Three main steering angles were selected at 5° intervals [−5°, 0°, 5°], with each main angle composed of three steering angles separated by 1.5° intervals. Raw radiofrequency (RF) data was acquired automatically at time intervals selected according to the kinetic behavior of each experimental model. For gelatin experiments, including gelatin-TG experiments, raw RF data was acquired every 2 minutes. The total duration of cooling plus acquisition was fixed at 135 min for all groups. Therefore, samples cooled for 15, 40, or 75 min were imaged for 120, 95, or 60 min, respectively. For fibrin clot experiments, raw RF data was acquired every 15 seconds for a total of 30 minutes. For blood coagulation experiments, raw RF data was acquired every 30 seconds for a total of 90 minutes. For blood thrombolysis experiments, raw RF data was acquired every 15 minutes for a total of 5 hours.

### 5.3 DSI algorithm

The DSI framework used in the present study estimates spatially resolved slowness deviations from temporal ultrasound echo displacements between sequential B-mode images. The DSI algorithm was previously developed in our group for monitoring temperature-induced acoustic changes during thermal ablation ^[43]^, and was subsequently adapted for cryoablation ^[44]^.

RF data was beamformed and interpolated onto a 599 × 390 pixel grid, corresponding to a field of view of 59 mm in the axial z direction and 40 mm in the lateral x direction. Each image underwent Hilbert transformation, after which the subsets of plane waves acquired at 1.5° intervals were coherently compounded to produce three main steering angle images per acquisition, corresponding to [−5°, 0°, 5°]. This coherent compounding step was used to improve signal quality prior to pixel tracking.

Dense optical flow was used to calculate pixel-wise echo displacements between consecutive B-mode images for each main steering angle using Farnebäck’s method ^[66]^. Dense optical-flow-based methods have previously shown promising results for spatially resolved displacement estimation in ultrasound elastography ^[67]^. The three calculated displacements acquired at each time interval were downsampled by a factor of 5 in the lateral direction and 10 in the axial direction and stacked to form a data vector Δd.

This downsampled displacement vector, Δd, was related to the slowness deviation map, Δσ, through the forward model:

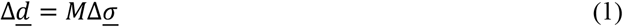

where Δ*d* represents the measured displacement vector, *M* is the forward model describing the contribution of local slowness to the observed displacement through angle-independent line integrals, and Δ*σ* is the spatial slowness deviation map.

Optimal slowness deviation maps, 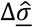, were estimated by solving a Tikhonov-regularized least-squares problem with L2 spatial-gradient regularization in the axial, lateral and diagonal directions:

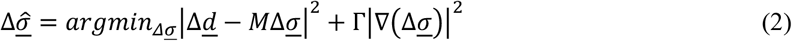

where the regularization term enforces spatial smoothness of the reconstructed slowness deviation map. In practice, the solution was implemented using a precomputed pseudoinverse operator, M^†^. The regularization parameters were chosen as Γ*x* = 10, Γz = Γxz = 1, corresponding to smoothness constraints in the lateral, axial, and diagonal directions, respectively, with increased smoothing in the lateral direction.

The resulting slowness deviation maps, 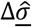, were subsequently upsampled to the original image dimensions. Because displacement estimation was performed sequentially and the reconstructed frame-wise slowness deviations were accumulated over time, the final maps represented cumulative slowness deviation relative to the initial state.

### 5.4 Quantitative analysis of speed-of-sound shift

To quantify temporal evolution in the reconstructed slowness deviation maps, representative slowness deviation values were extracted from each map. For clotting and gelation experiments, the maximum slowness deviation value was used. For lysis experiments, the minimum slowness deviation value was used. In practice, these slowness deviation values were consistently localized within the cylindrical inclusion. The SoS shift metric was calculated as the inverse of the representative slowness deviation value.

Because clotting/gelation and lysis produced opposite directions of apparent speckle motion in the B-mode images, the sign of the projected pixel-displacement field was defined according to the experimental condition prior to solving the inverse problem. This ensured consistency of the reconstruction framework across conditions.

### 5.5 Gelation time estimation by *τ*-based analysis

For each experimental run, gelation time was estimated from the SoS shift curves using a *τ*-based analysis. For each curve, the baseline plateau, C, was estimated as the median of the tail of the signal. The initial amplitude was defined as *A*_0_ = *y*(*t*_0_) − *C*, where *y*(*t*_0_) is the first value of the analyzed SoS shift curve. The characteristic time, *τ*, was defined as the first time point at which the curve reached the value *A* + *A*_0_ *e*^-1^. Prior to threshold detection, the curve was interpolated using shape-preserving piecewise cubic Hermite interpolation (PCHIP). The first interpolated crossing of the target threshold was used to determine *τ*. Unless otherwise stated, a value of 3τ, corresponding to approximately 95% of the plateau response, was used as the representative gelation or lysis endpoint throughout the study.

### 5.6 Statistics

Statistical analyses were conducted with the Prism10 software (GraphPad Software Inc.). Data is presented as mean ± SD. Each experimental group consisted of *n* = 3 independent experiments. The specific statistical tests used for each analysis are indicated in the corresponding figure captions. P values below 0.05 were considered significant.

### Supporting Information

Supporting Information is available from the Wiley Online Library or from the author.

## Acknowledgments

This work was supported in part by the Israel Science Foundation under Grant 192/22, in part by an ERC StG under Grant 101041118 (NanoBubbleBrain), the Israel Cancer Research Fund (grant number 1286686), in part by the Alrov center for Digital Medicine, and in part by the Nicholas and Elizabeth Slezak Super Center for Cardiac Research and Biomedical Engineering at Tel Aviv University.

Generative AI tools (ChatGPT, OpenAI, Claude, Anthropic and Gemini, Google) were used to assist with language editing and manuscript preparation. The authors reviewed and take full responsibility for all content.

## Conflict of Interest

The authors declare no conflict of interest.

## Author Contributions

S.G. designed and performed the research, conducted experiments, analyzed the data, and wrote the manuscript. T.G. guided and advised the analysis process. T.I. guided, advised, secured funding, and designed the research and wrote the manuscript. All authors reviewed the manuscript.

## Data Availability Statement

The data that support the findings of this study are available upon reasonable request from the authors. The code itself is patented and cannot be made publicly available.

## Notes

### Competing Interest Statement

The authors have declared no competing interest.

