## Supporting Information for "Ultrasound-Based Spatiotemporal Monitoring of Coagulation and Thrombolysis via Speed-of-Sound Shift Imaging"

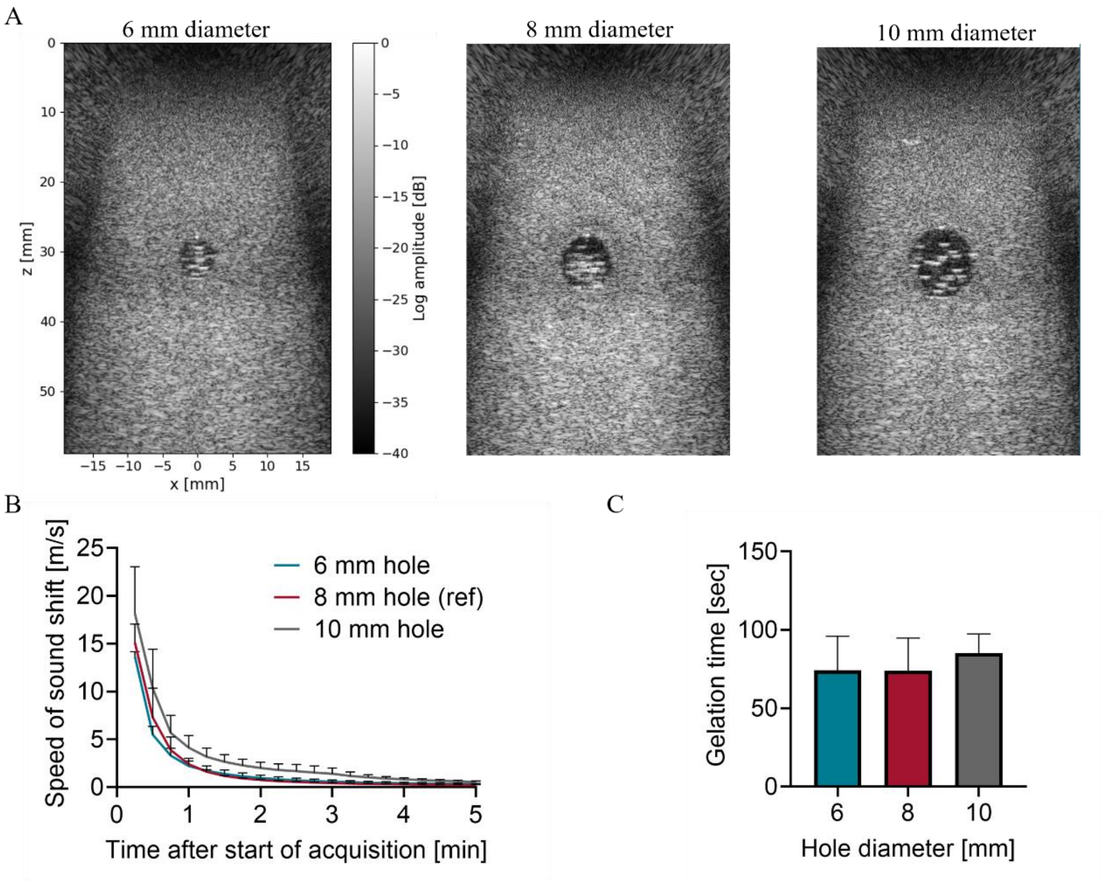


**Figure S1.** Effects of inclusion diameter on SoS shift and gelation time in fibrin clots. Fibrin clots were prepared with 2.5 mg $mL^{-1}$ fibrinogen, 0.65 U $mL^{-1}$ thrombin and 5 mM $CaCl_2$. Cylindrical inclusions of varying diameters were positioned 3 cm from the transducer and centered in the middle of the imaging plane. (A) B-mode images of agarose phantoms containing fibrin clots with 6 mm, 8 mm and 10 mm diameter inclusions (left to right). All B-mode images in (A) are displayed using the same field of view and the same dB scale (shown only for the left panel for clarity). (B) SoS shift as a function of time after the start of acquisitions in fibrin clots with different inclusion diameters. Inclusion diameter had no significant effect on SoS shift (Two-Way ANOVA with Tukey's multiple comparisons test) (C) Gelation times (3τ) for samples with different inclusion diameters, showing no significant differences (1-way ANOVA with Tukey's multiple comparisons test). Adjusted p values were *$p < 0.05$, **$p < 0.01$, ***$p < 0.001$ and ****$p < 0.0001$. N = 3. All data are plotted as the mean ± SD.

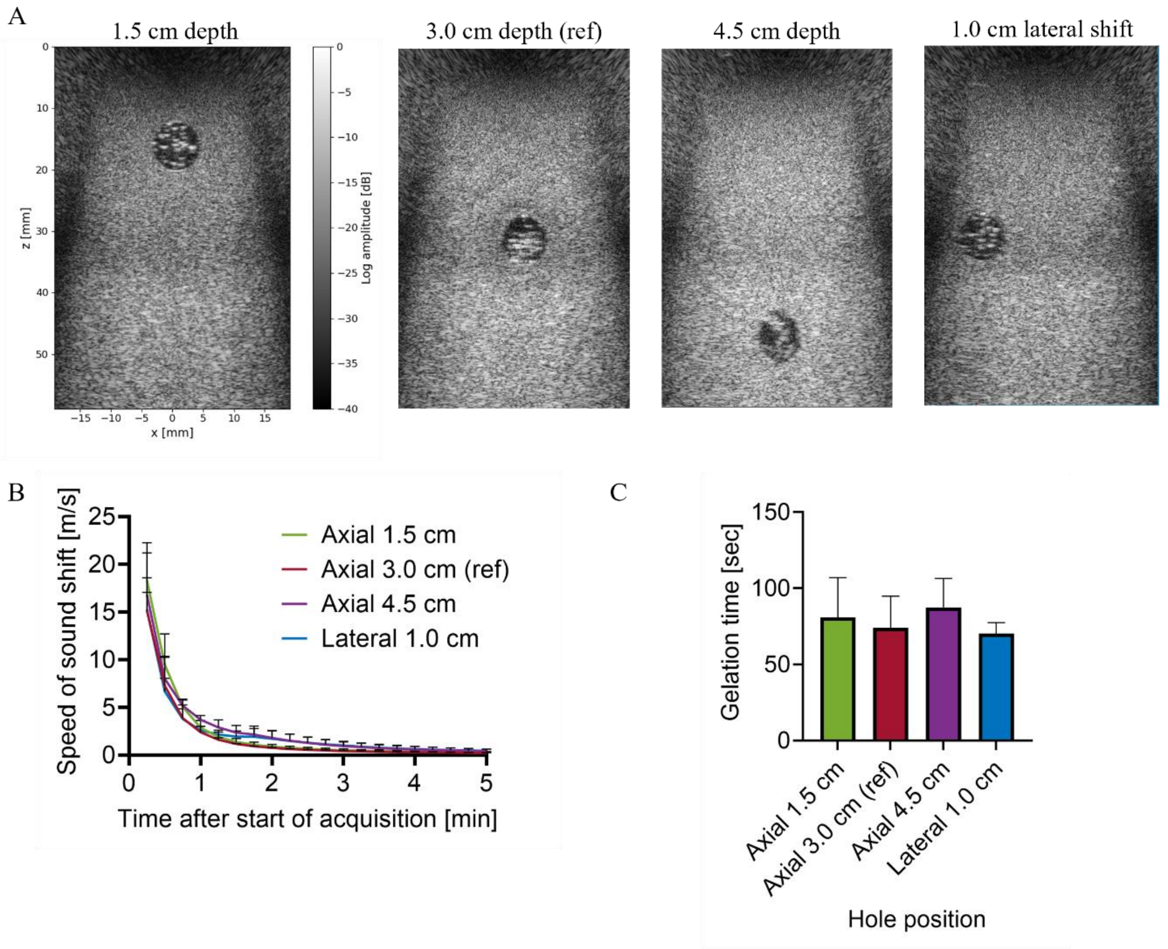


**Figure S2.** Effects of inclusion position on SoS shift and gelation time in fibrin clots. Fibrin clots were prepared with 2.5 mg $mL^{-1}$ fibrinogen, 0.65 U $mL^{-1}$ thrombin and 5 mM $CaCl_2$. Cylindrical inclusions with 8 mm diameter were positioned in different locations relative to the transducer. (A) B-mode images of agarose phantoms containing fibrin clots in inclusions in different positions, shown left to right: 1.5 cm depth, 3.0 cm depth (reference), 4.5 cm depth, and 3.0 cm depth with a 1.0 cm lateral shift. All B-mode images in (A) are displayed using the same field of view and the same dB scale (shown only for the left panel for clarity). (B) SoS shift as a function of time after the start of acquisitions in fibrin clots with different inclusion positions. inclusion position had no significant effect on SoS shift (Two-Way ANOVA with Tukey's multiple comparisons test). (C) Gelation times (3τ) for samples with different inclusion placements, showing no significant differences (1-way ANOVA with Tukey's multiple comparisons test). Adjusted p values were *$p < 0.05$,**$p < 0.01$, ***$p < 0.001$ and ****$p < 0.0001$. N = 3. All data are plotted as the mean ± SD.

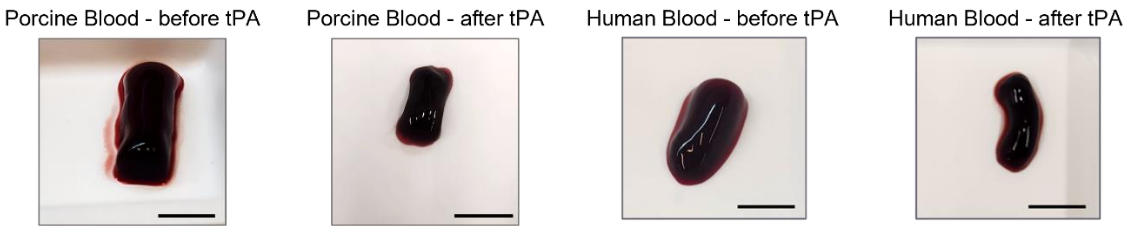


**Figure S3.** Representative photographs of porcine and human whole-blood clots before and after tPA-mediated thrombolysis. Clots were allowed to form following recalcification and were exposed to 6 µg $mL^{-1}$ tPA after 20 min of clotting. In both species, thrombolysis resulted in a visible reduction in clot size compared with the corresponding pre-treatment state. Scale bars = 1 cm.

**Table S1.** Effect of inclusion diameter on SoS shift and gelation time in fibrin clots. Values are mean ± SD (N = 3 per condition).

| Inclusion diameter [mm] | Distance from transducer [cm] | Lateral shift from image center [cm] | Initial SoS shift [m $s^{-1}$] | $\tau$ [sec] | 3$\tau$ [sec] |
|---|---|---|---|---|---|
| 6 | 3 | 0 | 13.70±0.37 | 32.06±3.62 | 66.17±10.85 |
| 8 | 3 | 0 | 15.14±1.57 | 34.73±7.26 | 74.20±21.78 |
| 10 | 3 | 0 | 18.17±3.98 | 38.41±4.05 | 85.23±12.16 |

**Table S2.** Effect of inclusion position on SoS shift and gelation time in fibrin clots. Values are mean ± SD (N = 3 per condition).

| Inclusion diameter [mm] | Distance from transducer [cm] | Lateral shift from image center [cm] | Initial SoS shift [m $s^{-1}$] | $\tau$[sec] | 3$\tau$ [sec] |
|---|---|---|---|---|---|
| 8 | 1.5 | 0 | 18.34±2.33 | 37.07±9.20 | 81.22±27.59 |
| 8 | 3 | 0 | 15.14±1.57 | 34.73±7.26 | 74.20±21.78 |
| 8 | 4.5 | 0 | 16.80±4.46 | 39.41±6.68 | 88.24±20.04 |
| 8 | 3 | 1 | 15.28±2.69 | 33.39±2.53 | 70.18±7.58 |
